# High-Resolution Characterization of rAAV Genomes for Vector Quality Using Long-Read Sequencing

**DOI:** 10.1101/2025.10.05.680573

**Authors:** Chao-Jung Wu, Hui-wen Liu, Thomas Carlile, Dongdong Lin, Baohong Zhang, Wei Zhang, Abdoulaye Baniré Diallo

**Affiliations:** Computer Science, UQAM, Montréal, Canada; Technical Development, Biogen, Cambridge, USA; Computational Biology & Genomics, Biogen, Cambridge, USA

**Keywords:** rAAV genotyping, residual DNA, long-read sequencing, PacBio, Nanopore, gene therapy vector quality

## Abstract

Recombinant adeno-associated virus (rAAV)-mediated gene therapy has been applied for human diseases. However, the rAAV capsids contain heterogeneous mixtures of full-length and truncated genomes and, depending on the manufacturing process, residual host cell and plasmid DNA. Therefore, a method is needed to characterize the encapsidated DNA of rAAV in order to support process development and batch release. The emerging long-read sequencing (LRS) has achieved AAV single-genome resolution. Here we propose a Python-based LRS profiling framework to classify and quantitate residual DNA species in rAAV products. We designed a reference that contains universal genetic components that are commonly used in rAAV production, including AmpR, KanR, Rep and Cap genes along with HPV18, Ad5 and hg38 genomes. We accessed the impurities of rAAV production from public and in-house LRS datasets. Analyzing the lambda fragments supplemented in these datasets showed that sequencing introduced size biases, which couldn’t be fully corrected by regression but is improvable within library preparation. Functional potential of impurities were assessed through indicators derived from long-read alignments, which enabled us to quantitatively compare impurities between manufacturing batches. We demonstrated that LRS provides informative metrics for rAAV production and can facilitate process development to ensure therapeutic product safety and quality.

## 1 Introduction

Adeno-associated virus (AAV), first discovered during adenovirus preparation in the 1960s, is a non-enveloped virus with a genome of single-stranded DNA (ssDNA) at 4.7 kb [Atchison et al., 1965; Hoggan et al., 1966]. Since then, tremendous discovery has been made and now recombinant AAV (rAAV) has become one of the most promising tools for delivering therapeutic human genes, given its lack of pathogenicity, target specificity, and long-term efficacy in multiple clinical trials and products [Hermonat and Muzyczka, 1984; Naso et al., 2017]. The typical structure of a rAAV cargo vector contains the gene-of-interest (GOI) around 4 kb in between the 5’-ITR and the 3’-ITR (inverted terminal repeats). Current manufacturing platforms to generate rAAV for clinical use include transient expression in human HEK293 cell line with plasmids, one carrying cargo vector, one carrying the AAV Rep/Cap genes and one carrying helper genes from adeno or herpes viruses; adenovirus-infected producer HeLa cell line; and baculovirus-infected insect Sf9 producing system [Naso et al., 2017]. AAV accumulates in both media and cellular lysate from producer cells. An AAV production will contain not only cellular debris but also heterogeneous populations of rAAV particles that range from containing the intact cargo (full capsides) to those without the ITR-flanked transgene (empty capsids) and populations of various sizes of the genome in between. Full capsids can be efficiently separated from empty capsids using anion exchange chromatography after initial AAV purification by heparin or AAV-specific affinity columns during Good Manufacturing Practice (GMP)-compliant commercial manufacturing [Naso et al., 2017; Potter et al., 2014]. However, one of the big challenges for rAAV manufacturing is the copurification of residual DNA. AAV can package various amounts of host cell DNA, AAV genes, helper plasmid DNA and selection markers (e.g. KanR or AmpR) along with the cargo DNA. Previous studies have shown the presence of heterogeneous genomic populations composed of truncated transgenes and contaminating sequences [Chadeuf et al., 2005; Tai et al., 2018]. These residual DNA are considered as impurities and more clinical experience is required to evaluate their adverse effects on product quality and safety [Wright, 2008]. Studies have shown that by choosing and modifying manufacturing platforms, the impurity profiles change [Tran et al., 2020, 2022]. Therefore, methods to assess product purity and integrity are essential for the optimization of manufacturing platforms. Current assessment methods for purity include electrophoresis such as alkaline agarose gel and capillary electrophoresis; genome identity can be verified by target-based approaches such as restriction fragment length polymorphism, quantitative polymerase chain reaction (qPCR), droplet digital PCR (ddPCR), and Sanger sequencing. These quality control methods (QC) are easy to implement and provide useful readouts at relatively low cost and short turn-around time. However, electrophoresis-based methods are limited by their low resolution. On the other hand, while target based methods are sensitive, accurate, and can quantify vector genome copy, whether there are rearrangements or unknown contaminating sequences in the AAV production batch are yet to be answered.

The next generation sequencing, including short-read sequencing (SRS) and the emerging long-read sequencing (LRS), is a powerful tool and could offer a solution [Oikonomopoulos et al., 2020]. While SRS gives valuable information regarding nucleotide variants and the extent of contaminating sequences, it poses limitations for assessment of full-length cargo because fragmentation of DNA input is required prior to sequencing library construction [Lecomte et al., 2015]. All next-generation sequencing platforms experience some degree of base-calling errors. However, advancements in library preparation methodologies have significantly reduced the error rates associated with long-read sequencing over the years [Oikonomopoulos et al., 2020]. For genotyping purposes, error rates below 10% are generally considered acceptable. In addition, GC bias of SRS results in low coverage of sequence alignment in GC-rich sequences and is known to aggravate genome assembly [Chen et al., 2013]. Sequencing through the ITR regions is also challenging due to its strong secondary structure [Tran et al., 2020].

Recently, studies based on LRS revealed the presence of truncated genomes and contaminating sequences fused with transgenes in the defective cargo [Tai et al., 2018]. The source of residual DNA is from packaging components such as AmpR, KanR, Rep and Cap genes along with HPV18, Ad5 and human genomes [Lecomte et al., 2015; Tai et al., 2018; Tran et al., 2020], depending on the manufacturing process [Dobrowsky et al., 2021]. These unintended sequences raise concerns regarding potential long-term expression in patient tissues, warranting close monitoring and control. Long-read sequencing has helped clarify the heterogeneity and complexity of residual DNA in rAAV genomes. To quantify this residual DNA, we developed a Python-based software for characterizing these sequences in purified rAAV products using a predefined reference residual DNA genome. ArcticFox accepts FASTQ data from long-read platforms, providing quantitative insights into impurities for cross-batch comparison using metrics derived from sequence alignments.

## 2 Materials and Methods

### 2.1 ArcticFox overview

ArcticFox includes modules specifically designed for the quantitative profiling of rAAV genome populations, as illustrated in Figure 2. It is compatible with sequencing data generated from major long-read platforms, such as PacBio and Oxford Nanopore. The LRS pipeline for assessing functional impurities in rAAV comprises three analytical modules integrated into the Python-based software ArcticFox: Dataset Validation, Genotyping and HG Analysis. ArcticFox is available as an open source project in GitHub in the following link https://github.com/bioinfoUQAM/ArcticFox

**Figure 1.**
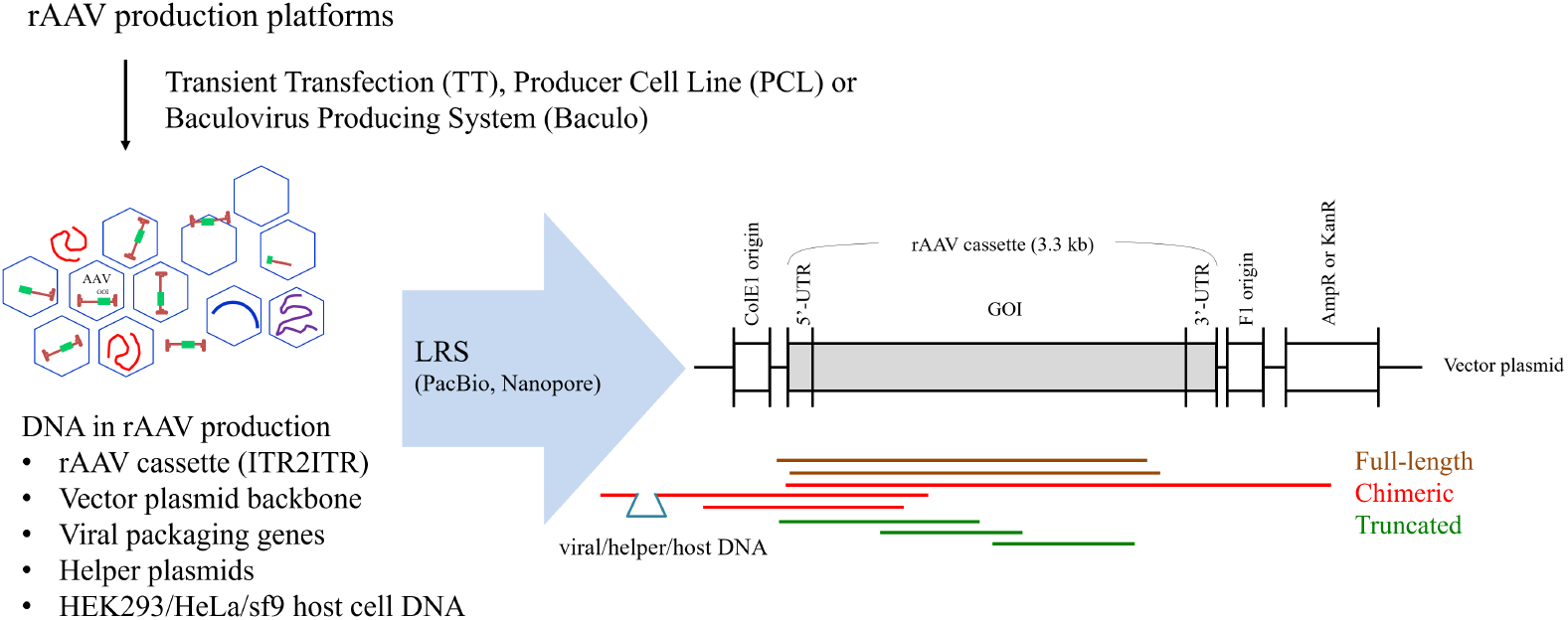
rAAV profiling by long-read sequencing. The rAAV capsids contain heterogeneous mixtures of full-length and truncated genomes and residual host cell and plasmid DNA, depending on the manufacturing process [Naso et al., 2017]. The emerging long-read sequencing (LRS) has AAV single-genome resolution [Tai et al., 2018].

**Figure 2.**
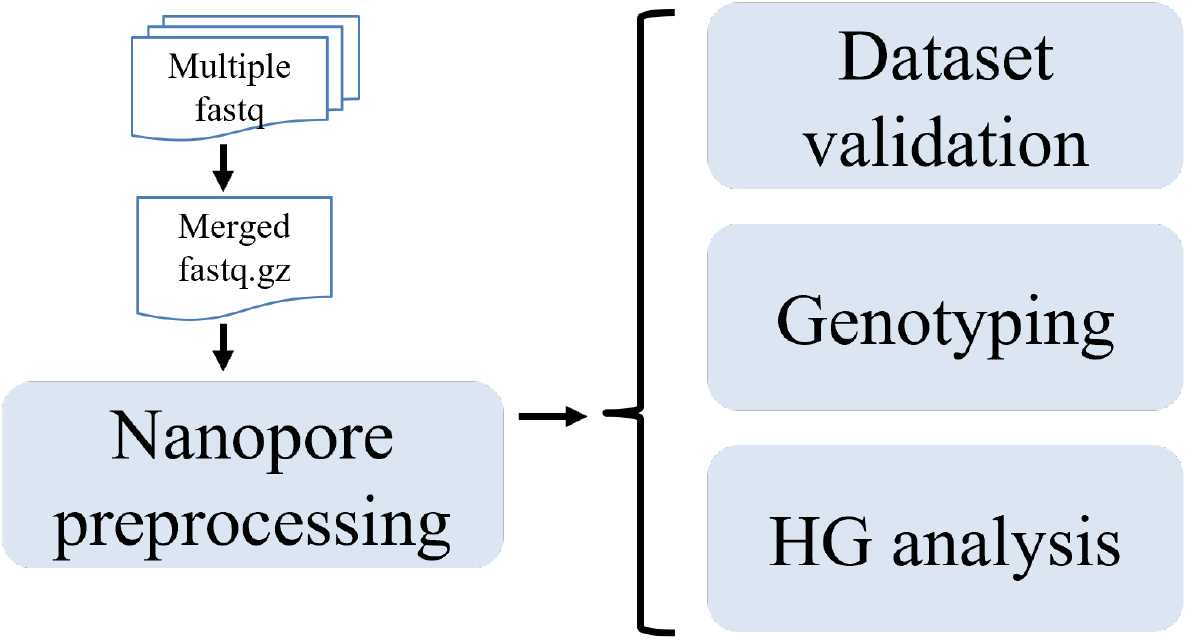
LRS pipeline for assessing functional impurities in rAAV, including three analytic modules embedded in the python-based software ArcticFox: Dataset validation to validate usable dataset, analyze lambda fragments and report picking up by template length and reading through by template length. Genotyping to analyze residual DNA based on several pre-defined chromosomes, reporting output metrics. HG analysis to analyze residual DNA coming from host genome hg38, reporting output metrics.

### 2.2 Datasets

To evaluate the consistency and utility of long-read sequencing for profiling rAAV genome populations, 18 LRS datasets from 4 rAAV production experiments were collected (Table 1). These datasets represent various rAAV production systems and sequencing data from PacBio and Nanopore platforms. In-house batches, Sample01 and Sample05, were prepared using different AAV production methods. Lambda fragments were added to the encapsidated AAV genomic DNA, followed by library preparation for Nanopore and PacBio sequencing. Selected public PacBio datasets were retrieved from the NCBI Sequence Read Archive. Lambda DNA readouts analyzed template length bias in sequencing procedures, and the experiments demonstrated the ability to compare genetic components using ArcticFox’s default reference-genome settings.

**Table 1.** LRS datasets of rAAV productions from public and in-house sources, including (a) inhouse, (b) SRA PRJNA383145 [Tai et al., 2018], (c) SRA PRJNA608034 [Tran et al., 2020], and (d) SRA PRJNA794197 [Tran et al., 2022], representing various rAAV production systems; and sequencing data obtained from PacBio and Nanopore long-read sequencers.

| Sample name | Transgene | Sequencer | Dataset source |
| --- | --- | --- | --- |
| Nanopore.Sample01 | BIIB-X | MinION Mk1B | a |
| Nanopore.Sample05 | BIIB-X | MinION Mk1B | a |
| PacBio.Sample01 | BIIB-X | PACBIO Sequel II | a |
| PacBio.Sample05 | BIIB-X | PACBIO Sequel II | a |
| pt2018_78 | EGFP | PacBio RS II | b |
| pt2018_79 | EGFP | PacBio RS II | b |
| pt2018_80 | EGFP | PacBio RS II | b |
| pt2018_81 | EGFP | PacBio RS II | b |
| pt2020_10 (Lib1) | Cas9 | PACBIO Sequel | c |
| pt2020_09 (Lib2) | Cas9 | PACBIO Sequel | c |
| pt2020_08 (Lib3) | Cas9 | PACBIO Sequel | c |
| pt2020_07 (Lib4) | Cas9 | PACBIO Sequel | c |
| pt2020_06 (Lib5) | mCherry | PACBIO Sequel | c |
| pt2020_05 (Lib6) | mCherry | PACBIO Sequel | c |
| pt2020_04 (Lib7) | EGFP | PACBIO Sequel | c |
| pt2020_03 (Lib8) | EGFP | PACBIO Sequel | c |
| pt2022_03 | EGFP | PACBIO Sequel II | d |
| pt2022_04 | EGFP | PACBIO Sequel II | d |

### 2.3 Design of reference genomes

The reference genomes supporting the analytic modules are illustrated in Figure 3. The first genome is for lambda calibration, containing 14 BstEII-digest fragments derived from the lambda genome. The second genome is for genotyping, containing 9 chromosomes: ITR2ITR, AmpR, KanR, Rep, Cap, HPV18, Ad5, lambda integral genome and backbone. The third genome is the human genome, containing 22 autosomal chromosomes, 2 sex chromosomes and 1 mitochondrial genome. The transgene in ITR2ITR is default Cas9 derived from pX602 (Addgene) or specific. The backbone is default pX602 removing transgene or specific. Lambda DNA-BstEII digest (New England Biolabs) is often supplemented as a sequencing control during library preparation.

**Figure 3.**
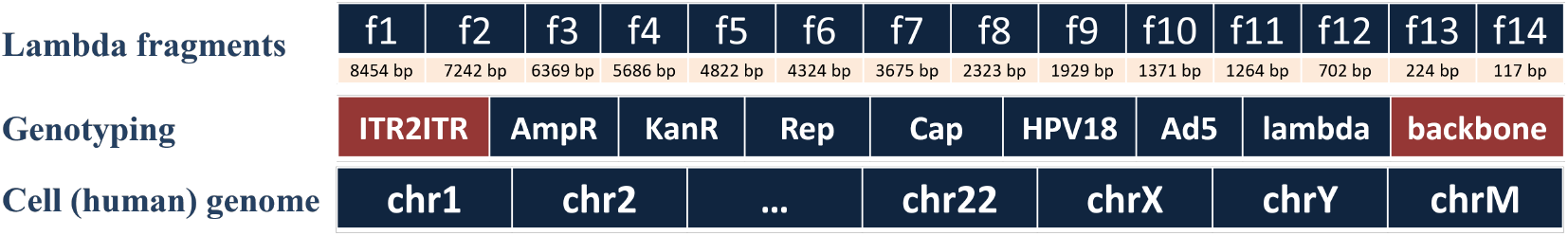
Design of reference genomes. The transgene in ITR2ITR is default Cas9 derived from pX602 (Addgene) or specific. The backbone is default pX602 removing transgene or specific. Lambda DNA-BstEII digest (New England Biolabs) is often supplemented as sequencing control during library preparation.

### 2.4 Correcting size biases of read abundance

We suggest that the size-biased read abundance could be modeled by regression as follows. Given *A*_*o*_: Observed read abundance, *L*_*t*_: Theoretical template length, *L*_*o*_: Observed read length and *R*_*o*_: Read abundance ratio (against baseline f6) obtained by analyzing lambda fragments, the corrected read abundance *A*_*c*_ in formula 3 can be obtained by determining formula 1 and 2, where formula 1 models the size bias regarding picking up template based on template length trained on lambda; and formula 2 models the size bias regarding complete reading through template based on template length trained on lambda. The model of formula 1 is a polynomial regression, and that of formula 2 is a linear regression. Formulas:

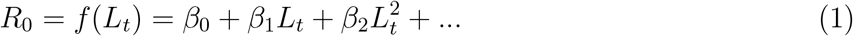

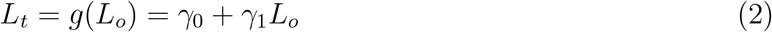

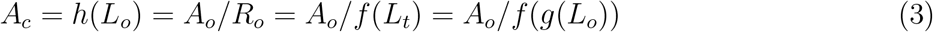

### 2.5 Functional quantification

#### Positivity rate

The positivity rate for each chromosome is defined as the percentage of reads in a batch aligned to that chromosome:

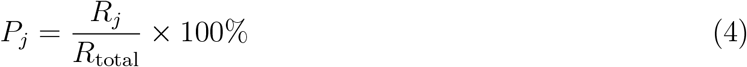

where *P*_*j*_ represents the positivity rate for chromosome *j, R*_*j*_ is the number of reads containing chromosome *j*, and *R*_total_ is the total number of reads in the library, excluding lambda reads, where *j* = { ITR2ITR, AmpR, KanR, Rep, Cap, HPV18, Ad5, backbone } . Take ITR2ITR as an example,

Positivity rate = reads containing ITR2ITR / total reads in library.

#### Intact rate

The intact rate for each chromosome is the percentage of reads that are quasi-full-length compared to all reads containing that chromosome:

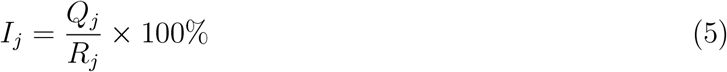

where *I*_*j*_ represents the intact rate for chromosome *j, Q*_*j*_ is the number of quasi-full-length reads of chromosome *j*, and *R*_*j*_ is the total number of reads containing chromosome *j*. Take ITR2ITR as an example,

Intact rate = reads are full length ITR2ITR / reads containing ITR2ITR; full-length threshold is 90%.

#### Mass percentage

Mass percentage for each chromosome is inferred by the proportion of nucleotides aligned to the chromosome relative to the total nucleotide count in the batch:

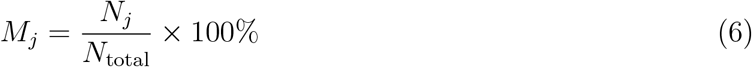

where *M*_*j*_ represents the mass percentage of chromosome *j, N*_*j*_ is the nucleotide count aligned to chromosome *j*, and *N*_total_ is the total nucleotide count in the library, excluding lambda nucleotides. Take ITR2ITR as an example, Mass percentage = nucleotides aligned to ITR2ITR / total nucleotides in library

## 3 Results

### 3.1 Size bias related to sequencing

Using known-length DNA fragments for spike-in analysis calibrates sequencer bias related to template length [Shirane et al., 2013; Castro-Wallace et al., 2017]. According to the Nanopore manufacturer’s guidelines, the spike-in procedure involves BstEII digestion of the lambda genome, producing 14 fragments (f1 to f14) of theoretically equal abundance. During sample preparation, lambda DNA fragments were added to each rAAV sample prior to sequencing. After sequencing, read abundance for each fragment was measured, and ratios were calculated against the 4324 bp fragment (Figure 4A). If sequencing is unbiased for picking up templates based on template lengths, these ratios should be close to 1. The completeness of reading for each DNA template is assessed by the full-length rate as the reading through rate, which should be 100% if there is no length bias (Figure 4B).

**Figure 4.**
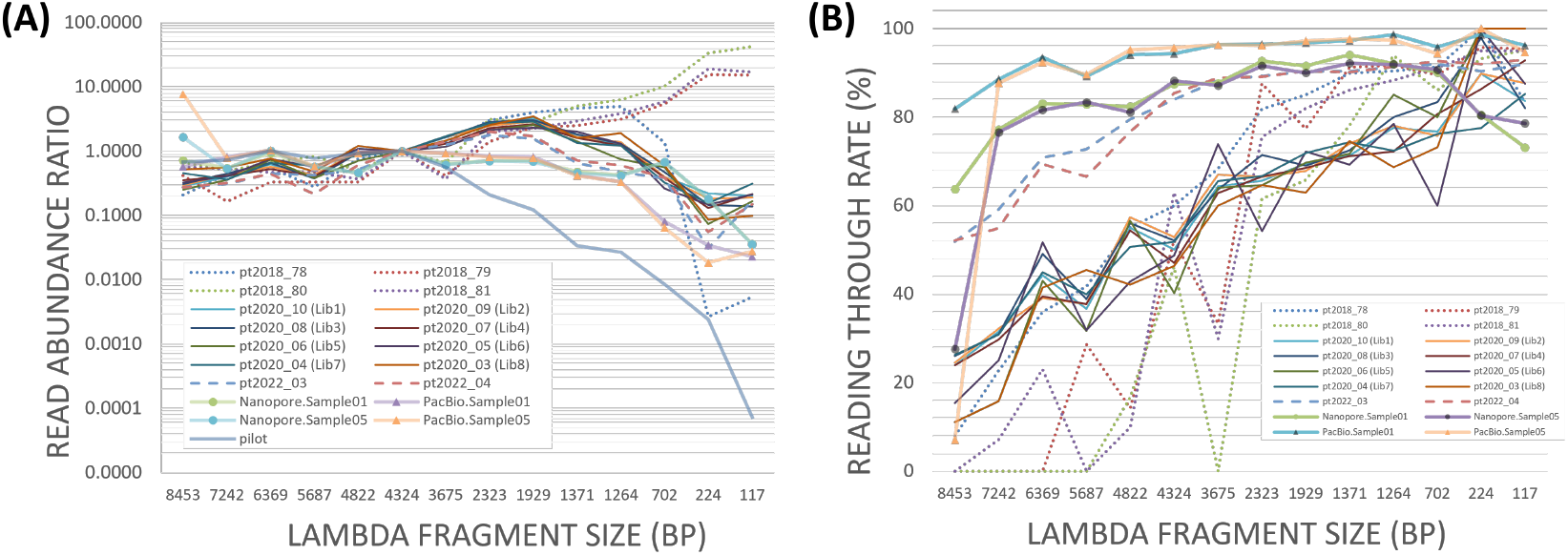
Size bias related to sequencing (A) regarding picking up template based on template length and (B) regarding complete reading through template based on template length. The BstEII-digestion of lambda genome gives rise to 14 fragments representing a range of template sizes that are theoretically of equal abundance ratio. After sequence identification, read abundance of each lambda fragment was measured and the abundance ratios were calculated against the fragment of 4324 bp. The theoretical reading through rate is 100%.

Picking up bias can be corrected if the ratios show a steady trend without fluctuations (Figure 5A2), otherwise negative ratio and overfitting occur in the polynomial regression model (Figure 5A1). For the regressions of reading through shown in Figure 5B, the values of the slope *γ*_1_ and the *R*^2^ are close to 1, which indicates that the major trend is *L*_*t*_ equals to *L*_*o*_ + *γ*_0_. However, reading through is challenging, as the square root of the mean squared error can reach several hundred base pairs, particularly when the theoretical length of observed small reads spans a wide range. Therefore, regressions have limited power to mathematically rescue the size biases and are only applicable for rescuing a steady trend of picking up bias. In-house and public datasets show that size biases are similar within experiments and improve over time (Figure 4), suggesting that library preparation protocols may reduce these biases.

**Figure 5.**
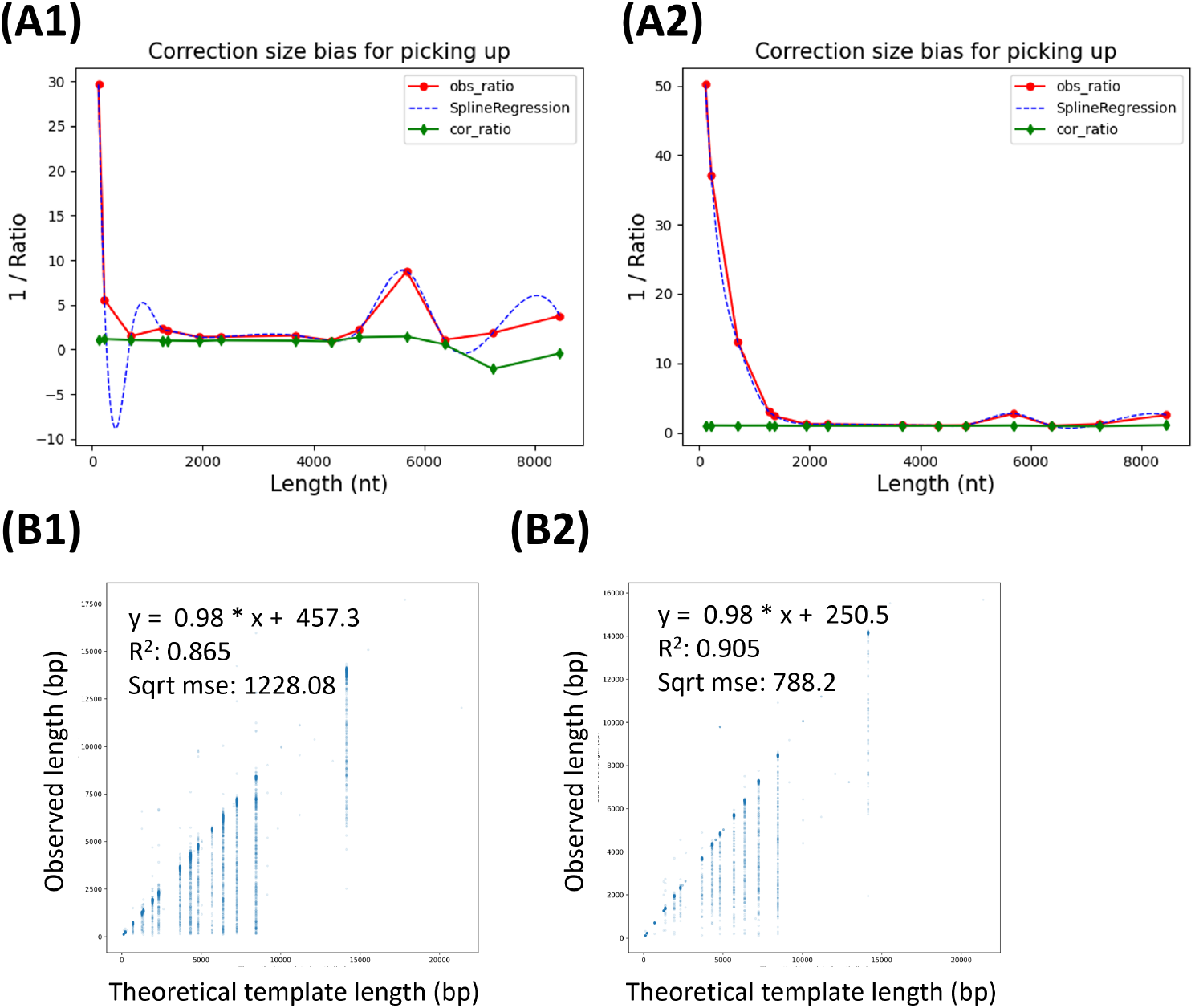
Correcting size biases of read abundance estimated from training on lambda in sample01 regarding (A) picking up and (B) reading through in (1) Nanopore and (2) PacBio. Picking up bias can be corrected if the ratios show a steady trend without fluctuations, otherwise negative ratio and overfitting occur in the polynomial regression model. Reading through is very difficult to correct, especially that the theoretical lengths of an observed small read have a wide range of possible values. Sqrt mse indicates on average how far (bp) the prediction of the linear regression model is from the theoretical length.

### 3.2 Multiple manufacturing batches

The changes of residual DNA profiles between batches and manufacturing processes can be captured by comparing the ArcticFox output metrics for the common genetic components Figure 6. Users can then select the most effective manufacturing process based on this comparative analysis, ensuring optimal quality and consistency in their rAAV products.

**Figure 6.**
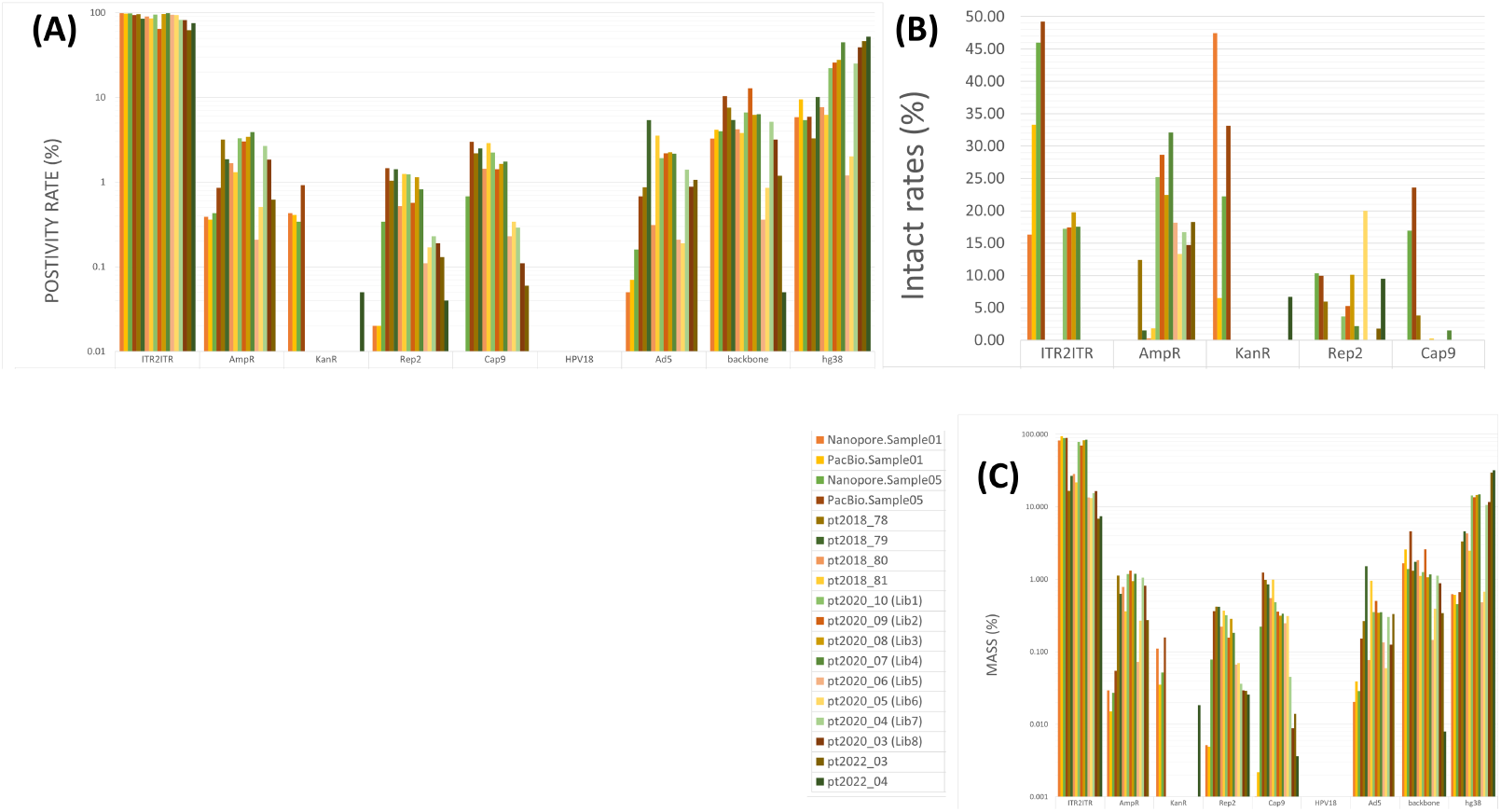
LRS sequencing provides several output metrics to evaluate functional potential of impurities, such as (A) Percentages of DNA populations based on reference genome positivity rates, (B) Intact rates of reference genomes, and (C) Mass percentages of reference genomes inferred from base count. These impurity metrics of the universal genetic components - AmpR, KanR, Rep and Cap genes and host genomes – allowed for quantitative comparison of impurities from different manufacturing batches; while the positivity rate can be compared if the ITR2ITR transgene between batches are the same.

## 4 Conclusion

LRS provides an opportunity to quick profile the drug substance before delivery or assess the outcome of a production procedure change or a vector design change. The universal reference genomes capture the genetic components that are commonly used in rAAV production. In addition to mass percentages of reference genomes, LRS provides the positivity rates of DNA populations and intact rates of reference genomes, which could not be obtained via short-read sequencing (SRS).

LRS is prone to size bias. Supplement of Lambda could be used to evaluate the quality of the library preparation and validate a usable dataset. The biases are similar within an experiment but different between experiments, indicating that the bias comes from library preparation rather than the sequencers. The size biases could be modeled by regression. However, overfitting may happen due to insufficient datapoints (e.g., 14 datapoints from standard fragments); and it is difficult to predict the theoretical lengths of an observed small read for modeling reading through. The correction power of regression is therefore limited. Improving library preparation is more intuitive than correcting the biases mathematically.

